# The representation of women as authors of submissions to ecology journals during the COVID-19 pandemic

**DOI:** 10.1101/2020.05.29.123455

**Authors:** Charles W. Fox

**Affiliations:** Department of Entomology, University of Kentucky, Lexington, Kentucky, United States 40546-0091

**Author notes:** Executive Editor of *Functional Ecology.

## Abstract

Observations made from papers submitted to preprint servers, and the speculation of editors on social media platforms, suggest that women are submitting fewer papers to scholarly journals than are men during the COVID-19 pandemic. Here I examine whether submissions by men and women to six ecology journals (all published by the British Ecological Society) have changed since the start of COVID disruptions. At these six ecology journals there is no evidence of a decline in the proportion of submissions that are authored by women (as either first or submitting author) since the start of the COVID-19 disruptions; the proportion of papers authored by women in the post-COVID period of 2020 has increased relative to the same period in 2019, and is higher than in the period pre-COVID in 2020. There is also no evidence of a change in the geographic pattern of submissions from across the globe.

---

Many studies have shown that female academics take on a disproportionately high share of family (Mason et al. 2013; Ecklund & Lincoln 2016; Sallee et al. 2016) and academic service (Guarino & Borden 2017) responsibilities, compared to their male professional colleagues, and that family responsibilities negatively affect the careers of women more than those of men (Mason et al. 2013). One consequence of the COVID-19 pandemic has been the closing of schools, child care facilities, and other public and private institutions that help manage children. Given the demonstrated disparities between men and women in their contributions to child care, it’s a sensible prediction that women will be more substantially affected by these disruptions caused by COVID-19 (Alon et al. 2020) and, in particular, that women will suffer more substantial drops in their scholarly productivity than will men during the pandemic.

Indeed, a few preliminary analyses have suggested this is the case. An analysis of submissions of economics papers to preprint servers (Shurchkov 2020) found a substantial (12-20%) decline in proportion of papers submitted by women in March and April 2020, compared to before the COVID-19 disruptions. A similar analysis of submissions to science preprint archives found that, overall, the rates of submissions of papers authored by women declined in March and April 2020, compared to both the preceding two months in 2020 and to the same two months (March and April) of 2019, though whether and to what extent there was a decline depended on the specific preprint server (Viglione 2020; Vincent-Lamarre et al. 2020). Some journal editors have also speculated on social media that they are observing fewer submissions authored by women post-COVID (discussed in Flaherty 2020 and Kitchener 2020), though the amount of data on which this speculation is based is limited. These observations made from papers submitted to preprint servers, and the speculation of editors on social media platforms, suggest that women are submitting fewer papers to scholarly journals than are men during the COVID-19 pandemic.

To test whether submissions by men and women to ecology journals have changed since the COVID pandemic became a significant influence on our lives and careers, I looked at submissions to six of the journals owned by the British Ecology Society (BES): *Functional Ecology* (FE), *Journal of Animal Ecology* (JAnim), *Journal of Applied Ecology* (JAppl), *Journal of Ecology* (JEcol), and *People and Nature* (PaN). I limited these analyses to submissions of standard research papers, submitted between 1 Jan 2019 and 21 May 2020 (the date the data were extracted); I’ve excluded reviews, commentaries and other non-standard or invited papers. I used the genderize.io database to infer author gender from author first names (using genderizeR, Wais 2016), as in previous studies (Fox et al. 2018, 2019; Fox & Paine 2019). Details on how data are extracted, genderized and analyzed are likewise presented in these previous publications by Charles Fox and colleagues.

## The representation of women as authors

Women were first authors on 40.8% of all submissions to these six ecology journals between 1 January 2019 and 21 May 2020 (averaged across journals), though this percentage varied among journals from a high of 51.0% to a low of 30.9% (Figure 1A). Women were similarly represented among submitting authors (the person who submitted the manuscript to ScholarOne Manuscripts); 39.1% of submitting authors were women (averaged across journals), though this varied among journals from a high of 51.2% to a low of 30.4% (Figure 1B).

**Figure 1.**
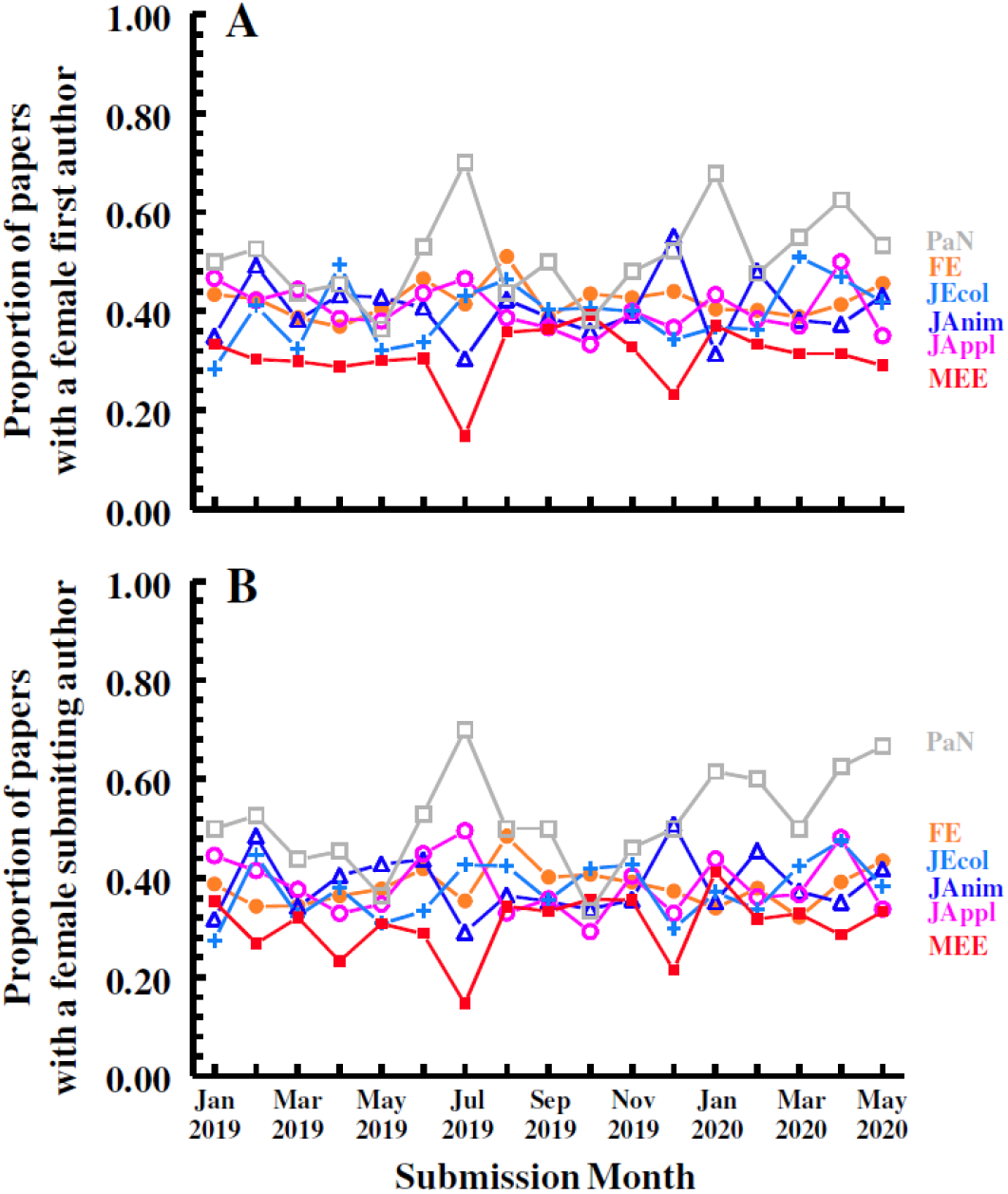
The proportion of papers for which the (A) first or (B) submitting author is a woman, for six of the journals published by the British Ecological Society. In total, these six journals received 7699 submissions from 1 Jan 2019 to 21 May 2020, ranging from a low of 302 submissions for *People and Nature* (PaN) to a high of 1869 submissions to *Functional Ecology* (FE).

**Figure 2.**
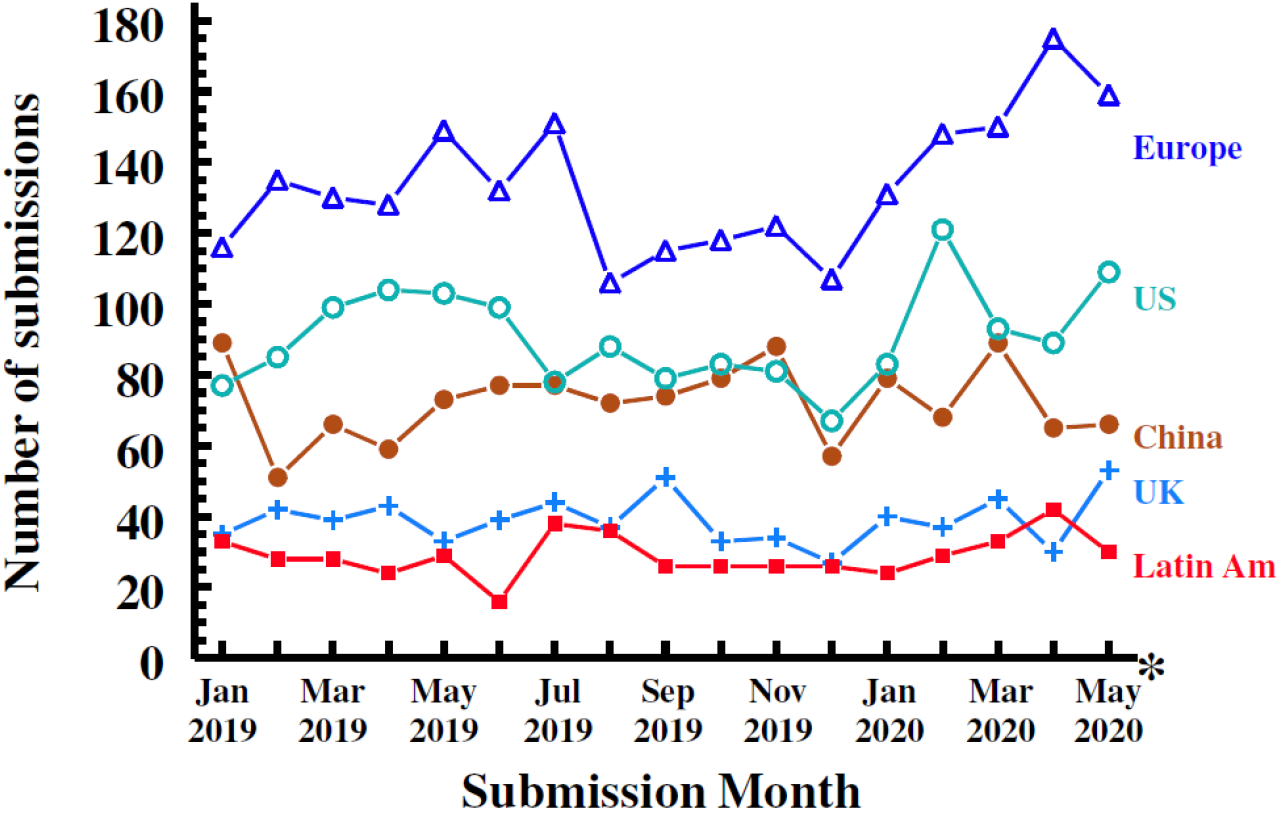
Monthly number of submissions, cumulative across the five journals, from five major regions of the world, mainland Europe (△), the United States (US; ○), China (●), the United Kingdom (UK; +), and Latin America (North and South America, excluding the US and Canada; ■). Because I only have data through 21 May 2020, submissions for May 2020 (*) are extrapolated from the average daily number of submissions over the first two-thirds of the month.

To test whether the representation of women as authors of ecology papers has changed since the start of the pandemic disruptions and stay-at-home orders, I had to establish a date that we can consider to be the start of the disruptions. Any chosen date will necessarily be imprecise since every country, and even regions within countries, instituted stay-at-home orders and closed their schools and universities on different dates. I chose 15 March 2020 as the start of the post-COVID disruption period for these analyses because this is roughly the middle of the 2-3 week period during which the majority of United States universities closed to students and researchers. Countries, and even regions or universities within countries, vary substantially in the degree to which they exclude researchers from their offices and labs, and this variance is not captured in these analyses.

First, I compared the proportion of papers with female authors during the period 15 March 2019 to 21 May 2020 with the same period (15 March to 21 May) in 2019. Contrary to the observations from some preprint servers and the speculation of editors of some journals (citations above), the proportion of both first and submitting authors that are women actually trends towards being higher in the post-COVID period of 2020 than in the same period of 2019. 42.7% of all submitted manuscripts had a female first author in the post-COVID period of 2020, compared to just 37.5% of all papers in the same period of 2019 (logistic regression comparing years with journal as a random effect; *X*_1_ = 3.1, *P* = 0.08). Similarly, 42.6% of papers had a female submitting author in 2020, compared to just 35.4% in 2019, a statistically significant increase in the proportion of papers submitted by women (*X*_1_ = 5.2, *P* < 0.001).

The above analysis is instructive but is possibly biased by the observation that the proportion of papers authored by women has been steadily increasing year-to-year at these six ecology journals (Fox et al. 2018). I thus also compared submissions in the 67 days pre-15 March 2020 with submission in the 67 days from 15 March to 21 May 2020. As with the previous analysis, I found no evidence that the proportion of first or submitting authors that are women has declined; 42.4% of papers had female first authors pre-COVID, compared to 42.7% post-COVID (*X*_1_ = 0.02, *P* = 0.89), and 40.6% of submitting authors were women pre-COVID, compared to 42.6% post-COVID (*X*_1_ = 0.14, *P* = 0.71).

In previous studies of submissions to ecology journals (Fox et al. 2016, 2018), we have observed that the first author is less likely to be the person that submits their manuscript (and serve as corresponding author) if they are female than if they are male. In the current dataset (1 January 2019 to 21 May 2020), 83% of papers were submitted by the first author. Female first authors were slightly less likely than male first authors to be the submitting author (logistic regression with journal as a random effect; *X*_1_ = 4.6, *P* = 0.03), though the effect is small (only 2%), much smaller than the ~9% difference reported in the much larger dataset of Fox et al. (2018). There was no evidence that the likelihood that a female first author served as submitting author differed pre- vs post-COVID, either comparing similar timeframes between years (*X*_1_, = 1.34, *P* = 0.24) or comparing pre- vs post-COVID in 2020 (*X*_1_ = 0.56, *P* = 0.45), though the dataset is quite small and thus has very low power for detecting interactions in a logistic regression.

The conclusion of these analyses is that, at these six ecology journals, I don’t see evidence that the submission of manuscripts by women have been more greatly affected than submissions by men by COVID-19 disruptions. It’s important to be clear, though, that this analysis only captures submissions during a short window of time, many of which may have been written before the COVID disruptions. We need to extend this analysis later this year before we can convincingly conclude that submissions by women are being impacted similarly to those by men.

## Number of submissions

The analysis above demonstrates that men and women are similarly represented in submissions to the BES journals both before and after the COVID disruptions. But that analysis does not consider whether total submissions to these six ecology journals (irrespective of author gender) have declined, remained robust, or even increased (since many researchers are barred from their labs and/or field sites and may have little else to do than write), during the COVID pandemic. I thus compared the number of submissions received by these journals post-COVID disruption (15 March 2020 to 21 May 2020) with those received during an identical period last year in 2019. Submissions to these journals were 7.6% greater (cumulative across journals) in 2020 than in 2019 (1072 submissions in 2019 versus 1153 in 2020). This overall increase remains positive (7.0%; from 1041 submissions in 2019 to 1113 submissions in 2020) after excluding *People and Nature*, the newest journal in the stable, which increased the most (29%) between the two years. A 7% increase is similar to the year-on-year increase in submissions experienced by these journals over the past few years, so not out of line from normal. That submissions were largely unaffected by the COVID disruptions is likewise suggested by a comparison of the number of submissions for the 67 days pre-March 15 and those during the 67 days from 15 March to 21 May 2020; these journals received a total of 1153 submissions (3% more papers) in the 67 days after the COVID disruption compared to the 67 days before the disruptions (1121 submissions).

The dataset post-COVID is inadequate to do a more detailed analysis of whether specific regions of the world are more greatly impacted by COVID disruptions. However, visual inspection of submission data broken down by five major submission regions (Figure 4) does not suggest any particular regions show unusual patterns of submission in March-May 2020, though patterns can be obscured by substantial among-month variability.

## Final thoughts

The closing of universities and other businesses, and the need to socially isolate to reduce transmission of COVID-19, have profoundly disrupted nearly every aspect of scholarly research. It’s inevitable that these disruptions will reduce the amount of scholarly research done until the pandemic slows and it’s a reasonable prediction that women will be more greatly impacted than men if they bear a greater share of the increased caretaker workload caused by the closing of our schools and child care facilities. However, contrary to some other preliminary analyses of submissions to preprint servers (Shurchkov 2020; Vincent-Lamarre et al. 2020) and the speculation of editors on social media (Flaherty 2020), we do not yet see evidence that there has been a decline in the number of submissions by women to the ecology journals published by the BES. We also do not see evidence of a change in the geographic pattern of submissions from across the globe, despite some areas being particularly heavily impacted by early pandemic disruptions. Of course, papers being submitted today are the result of work done months or years prior to submission, and so it may be too early to begin seeing the pandemic’s impacts; effects of COVID-caused disruptions may not become evident for months or years to come. We thus need to revisit the question later this year.

Although we do not yet see a differential in the effect of COVID-19 on submissions by men and women, the widespread closing of primary schools and child care facilities will have disproportionately large effects on some members of our community. It’s thus critical that academic institutions and scholarly societies develop the infrastructure - both procedures and policies - for addressing the various inequalities that have been and will continue to be created by this pandemic. Most importantly, these institutions need to support early career researchers - our students and recent graduates whose careers are most vulnerable to extended disruptions to their research and the resulting reduced scholarly productivity. Vincent-Lamarre and colleagues (2020) provide a list of suggestions for how best to support our vulnerable colleagues, many of which, we are pleased to see, have been adopted by leading universities.

## Acknowledgements

Thanks very much to the Assistant Editors of the six BES journals (Simon Hoggart, Jennifer Meyer, Alice Plane, Rhiannon Robins, Kirsty Scandrett and India Stephenson) for extracting these data from ScholarOne Manuscripts. Thanks also to Josiah Ritchey for running GenderizeR, and to Ruth Bryan, Emilie Aimé and Catherine Hill for helpful comments on earlier versions of this paper.

